# Egocentric anchoring-and-adjustment of social knowledge in the hippocampal formation

**DOI:** 10.1101/2024.09.13.612825

**Authors:** Marta Rodríguez Aramendía, Mariachiara Esposito, Raphael Kaplan

## Abstract

Recent work suggests the hippocampal formation(HF) assimilates relational social knowledge similar to how it transforms egocentric spatial cues into map-like representations. Yet whether hippocampal map-like representations of social knowledge still represent lingering egocentric biases is unclear. We test if a prominent egocentric bias involving an implicit reliance on self-knowledge when rating others, anchoring-and-adjustment, is present when the relative attributes of different social entities are assimilated by the HF. Participants provided likelihood ratings of partaking in everyday activities for themselves, fictitious individuals, and familiar social groups. Adapting a functional neuroimaging task from Kaplan and Friston, participants then learned a stranger’s preference for an activity relative to one of the fictitious individuals and inferred how the stranger’s preference related to the groups’ preferences. Egocentric anchoring-and-adjustment was present when participants rated the other entities. Isolating the neural representation of egocentric anchoring-and-adjustment when flexibly comparing different social entities, the HF and dorsomedial prefrontal cortex(dmPFC) represented group-self rating discrepancy. Furthermore, the HF also reflected how well group preferences were remembered, where memory for group preferences correlates with task performance. We found the HF selectively represented group identity over other learned entities, confirming that the HF was primarily engaged by social comparisons in a more ample frame of reference. Taken together, these results imply that self-knowledge influences how the HF assimilates map-like knowledge about others.

## Introduction

Learning via cognitive maps permits an organism to relate their current state to an internal model of the world [1]. Cognitive maps have inspired a better understanding of how neurons in the hippocampal formation(HF), including entorhinal cortex (EC), transform egocentric spatial cues into purely environment-centered allocentric coordinates supporting spatial navigation [2]. Growing evidence suggests the HF assimilates abstract knowledge similar to how it transforms spatial cues [3–6], where social learning tasks have been particularly successful in isolating this new role[7-11; see 12 for review]. In particular, Kaplan & Friston(2019) investigated social learning beyond well-studied self versus other comparisons by testing how a stranger’s preference was learned relative to one individual and then compared to other individuals. The transformation from learning preferences relative to one point of reference and relating that preference to others highlighted a role for the HF in social inferences in an absolute reference frame(i.e., flexibly comparing everyone in one’s social environment). Although comparisons were made on a 1D number line of preferences, the authors viewed an absolute reference frame as analogous to map-like allocentric reference frames. This analogy is inspired by the similarity between the paradigm’s nonlinear relative to absolute reference frame transformations of preferences and nonlinear egocentric-allocentric spatial coordinate transformations made during spatial navigation[13,14]. The extension of allocentric/absolute coding principles to social learning raises the possibility that spatial and social perspective-taking could rely on the same neural computations in the HF and beyond[15]. However, whether this form of hippocampal social learning is supported by an isometric map-like representation, or an integrated model of relational knowledge via more imprecise subjective mechanisms is unknown[16–18]. On one hand, firing fields of spatially-modulated hippocampal neurons resemble cartesian coordinates on a map[5,19] , yet the self often serves as a reference point when making memory-guided inferences[20,21]. Moreover, previous work involving social reference frame transformations used the self as one of the explicit reference points in all conditions[8]. Consequently, it is unclear if subjective biases might be present in HF map-like representations of abstract knowledge about others.

One well-characterized subjective bias that could occur during map-like learning is anchoring-and-adjustment, where an individual starts with an initial idea and incompletely shifts away from their initial starting point to make an inference[22]. In egocentric anchoring-and-adjustment, people commonly begin by recruiting self-knowledge and then adjust away from this self anchor to make inferences about others(e.g., you judge someone else’s preference for a food that you like to be closer to your own preference than it should be)[23,24]. Notably, the more divergent others’ attributes are from a participant’s, the more adjustment is needed and the longer it takes to make the inference [25,26]. These findings are consistent with the proposition that adjustment from self-generated anchors occurs serially over time[21]. Relating anchoring to neural processes, social anchoring biases, where dorsomedial prefrontal cortex(dmPFC) tracks the divergence between self and other preferences[27]. Although egocentric anchoring is well-described in social decision tasks[27], evidence of its presence in map-like knowledge representations of the social world is missing[8,15]. This dearth of evidence is due to egocentric anchoring being tested on self v. other comparisons. Such one to one comparisons don’t require the type of nonlinear transformations needed to simultaneously relate different people to each other and the world in a common reference frame[8,15], or transform the position of an egocentrically viewed spatial cue into a location in an allocentric map of the environment[13,14].

Clues about how self-knowledge implicitly informs map-like representations of abstract knowledge come from work by Kaplan & Friston(2019). The authors had participants learn a stranger’s preference for an everyday activity—relative to previously provided ratings for either themselves, a close friend, or a typical person on that same activity—and subsequently decide how the stranger’s preference relates to the other two individuals’ preferences. Highlighting how the metric nature of these abstract distances influenced behavior, they observed a steady increase in the quickness and accuracy of the participant’s responses when the stranger’s rating was closer to one option versus another. In parallel, they identified a HF region relating to the absolute distance between the stranger’s rating and the choice options differently depending on whether a self-comparison was present in the absolute reference frame[8]. Still, due to self preferences always being needed in the transformation and/or comparison process, it is unclear whether egocentric anchoring may occur during the task. By removing the explicit need to use self-knowledge during the inference process, while keeping everything else the same, we can measure the implicit representation of self preferences tied to other social entities during relative to absolute reference frame transformations.

Here, to investigate whether there is an implicit use of the self on social inferences in an absolute map-like reference frame, we adapted the Kaplan & Friston(2019) paradigm by collecting participants’ preference ratings on everyday activities(e.g., eat spicy food, read a book, cycle to work)for themselves, two fictitious individuals, and two societal groups (people from cities and rural areas) from 1 to 9 on a 0-10 (ranging from impossible to happening at this moment) scale to allow for strangers with higher or lower ratings than the entities in the subsequent fMRI task(Fig. 1A). To expedite learning of the fictitious individuals, we made the two individuals congruent with one of the two rated groups(a rural or urban person). Subsequently for the transformation phase that occurred while participants underwent functional magnetic resonance imaging(fMRI), participants inferred a stranger’s preference for an everyday activity—relative to previously provided ratings for one of the two individuals—and decided how a stranger’s preference relates to a medium preference(normal group) and a preference of one of the two groups. More specifically, participants needed to decide whether the stranger’s rating was closer to one of the two groups, or a normal group with a medium rating of 5(Fig. 1B). After deciding without any feedback, a jittered intertrial interval period(mean 2.5s) featuring a black fixation point overlaid on a gray background appeared on the screen before moving onto a new trial. After the fMRI task, participants provided ratings for all entities, including the self, on the same scenarios to assess consistency/memory for ratings.

**Figure 1.**
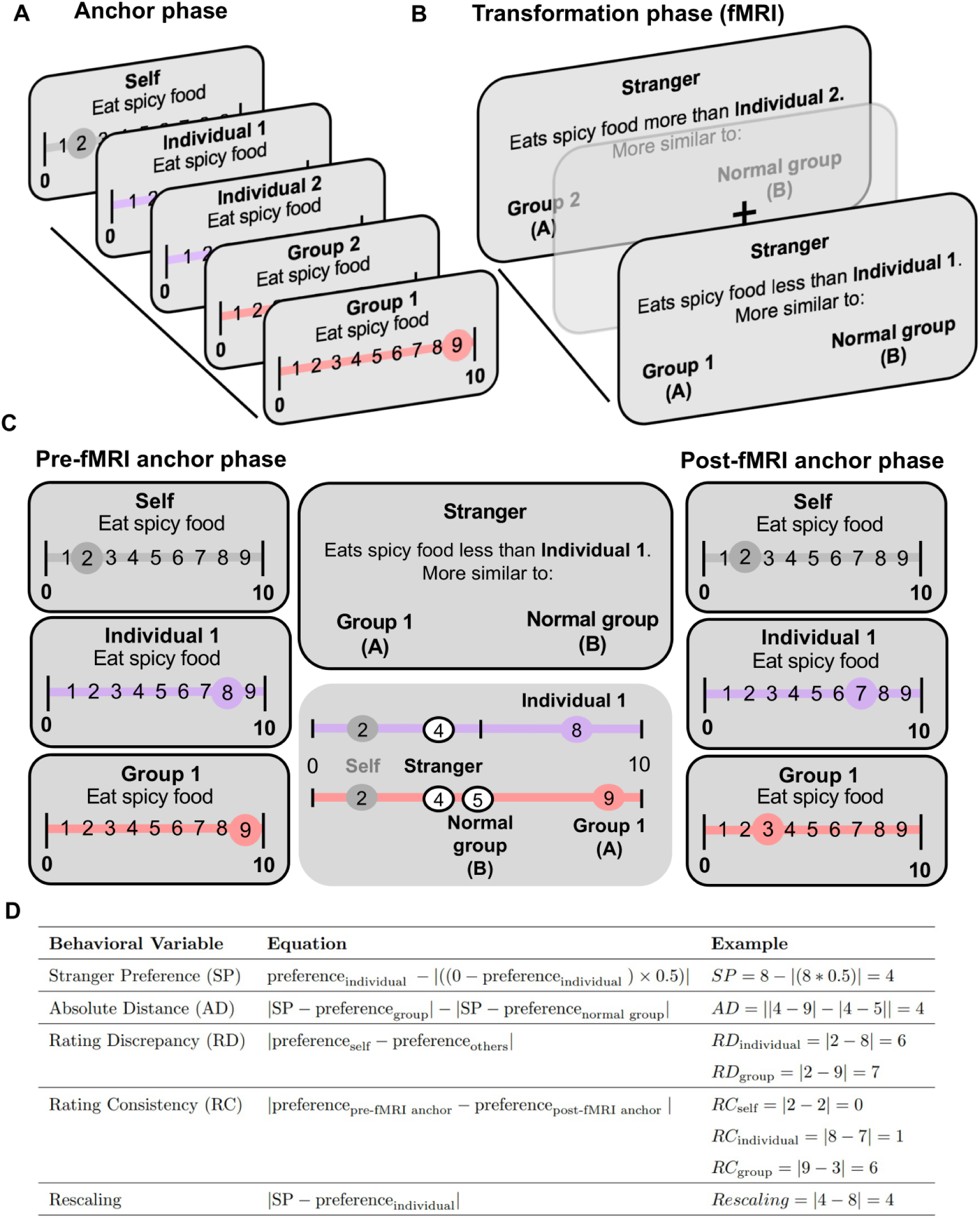
Experimental paradigm. (A) Anchor phase. Before and after fMRI scanning, participants provide a likelihood rating for themselves in different everyday scenarios, as well as confidence ratings about each preference. After reading a description of two different fictive individuals and being presented with two different social groups, participants rate the likelihood of these entities from 1-9(very unlikely to very likely) on a scale of 0 to 10 to partake in a given everyday scenario. (B) Transformation phase. During fMRI scanning, participants infer a stranger’s preference relative to one of the previously rated individuals in a particular scenario and in a two-alternative forced choice(2AFC) determine whether that stranger’s preference is more similar to a normal group(participants were informed beforehand that this group always has a preference rating of 5), or one of the two groups they previously rated(using the preference rating provided by the participant for that entity). The 2AFC was self-paced(mean=4.48s) and the intertrial interval duration lasted a mean of 2.5s(range 1-4s) over four runs. (C) Example fMRI trial where participantś pre- and post-fMRI task ratings for themselves(the rating in gray) and the other relevant entities(individuals in purple, groups in pink) are highlighted. Ratings for the self aren’t needed to accurately perform the fMRI task. (D) Table describes how the experimental variables of interest would be calculated for the given preference ratings in this example trial in C. Stranger preferences(SP) are only used to inform the other behavioral variables and aren’t analyzed in isolation in this study. Crucially, post-fMRI task ratings are only used for the Rating Consistency(RC) variables.

Including two different types of both individuals and groups ensured that the rated entities, the stranger, and the participant didn’t have overlapping ratings too often(see Methods for more details). Furthermore, separating the groups and individuals in different stages of a transformation trial allowed us to test what task representations were most important to the HF and dmPFC(Fig. 1C). Crucially, self ratings weren’t necessary to perform the transformation phase of the task and unlike the previous version of the task, no spatial number line cues of preferences were provided to remove potential spatial scaffolding effects that could enhance performance. This experimental design allows us to study how the absolute distance between the stranger’s rating and the ratings of the groups that are being compared with the stranger(absolute distance, AD), the differences between self and others’ preferences(rating discrepancy, RD, with either groups or individuals), the cognitive demand of how far the stranger’s preference is away from the individual’s rating(rescaling), and the memory for self and others’ preferences(rating consistency, RC, for groups or individuals) are represented during reference frame transformations of social knowledge(see Methods and Fig.1D).

The motivations behind our experimental design were threefold: First, to replicate both the egocentric anchoring-and-adjustment rating response time(RT) and accuracy findings in the social domain[25,26]; Second, to replicate the linear relationship between accuracy and the absolute distance between the stranger’s rating and the compared groups’ in the transformation phase[8]; Third, to isolate the behavioral and neural representation of egocentric anchor biases during reference frame transformations of social knowledge. We predicted that egocentric anchoring-and-adjustment would be present in map-like representations of social knowledge in the hippocampal-entorhinal system and dmPFC. We employed representational similarity analysis(RSA) here because we were interested if trial-level differences in self-other(group or individual) rating discrepancy, independent of preference type(i.e., food, transportation, music, etc.), would be represented in a common format in the aforementioned regions during our social inference task.

## Results

### Behavioral results

During the anchor phase, the average time that participants took to provide ratings of others was 3.12s(SEM=0.085). We ran a linear mixed effects model to test the influence of various factors on determining self versus other discrepancy for each participant. We observed that the anchor entity, individuals and groups, influenced how participants judged the target. A significant effect of entity was observed (*b*=0.077, t(19)=2.10, p=0.036), indicating that participants rated the preferences of individual entities closer to their own preferences than those of group targets. Examining the hypothesized positive linear relationship between RT and self versus other discrepancy, we found a significant effect of RT on self-other discrepancy scores (*b*=0.21, t(19)=2.18, p=0.030). This result suggests that increases in RT correspond to increases in the magnitude of correction away from a self-anchor—evidence of serial adjustment present in anchoring-and-adjustment(Fig.2A).

**Figure 2.**
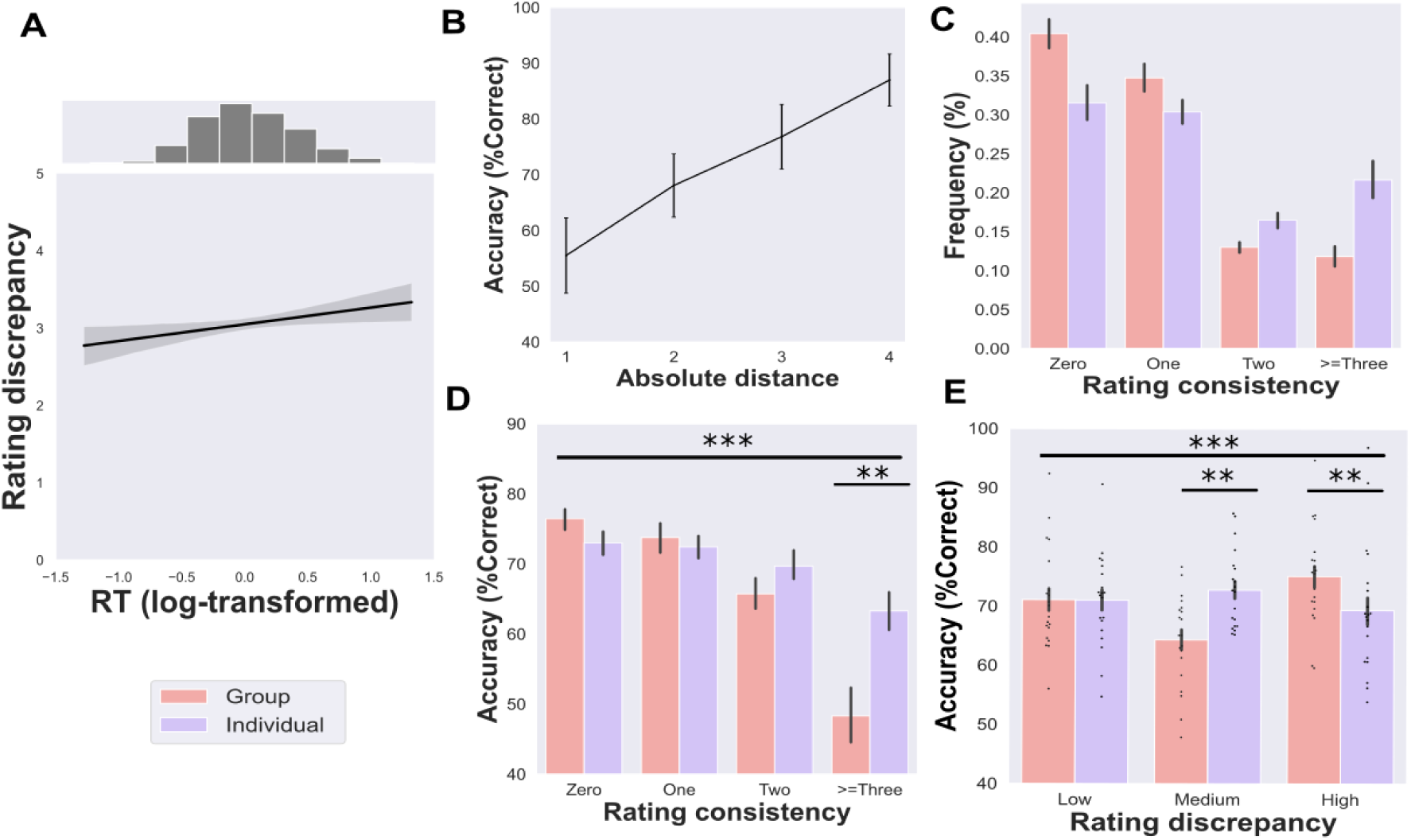
Behavioral results: (A) Positive relationship between self versus other entity rating discrepancy(RD) and reaction time(RT: centered by group mean). (B) Significant relationship between accuracy and absolute distance between the stranger’s rating and two choice options for each trial. (C) Distribution of group and individual rating consistency (RC). (D) Significant difference in accuracy for rating discrepancy for groups and individuals. (E) Significant differences between individuals and group rating discrepancy and accuracy. All error bars showing mean ± SEM with dots representing individual participants. Asterisks showing significant differences: *= p<.05, ** = p<0.01 , *** =p<0.001.

To test if serial adjustment from self-knowledge occurred exclusively when the target was an individual or a group, we explored whether the relationship between RT and self–other discrepancy depended on entity type(i.e., individuals or groups). There was no significant interaction of the target entity on RT (*b*=-0.07, t(19)=-0.68, p=0.49). These findings replicate prior social anchoring-and-adjustment studies[25,26], aligning with the idea that participants anchor on self-knowledge and serially adjust away from this anchor when evaluating both individuals and groups.

To study whether self-other discrepancy was consistent across the anchor phase, we calculated and compared pre-fMRI and post-fMRI RD using the ratings provided in each of the anchor phases(Fig 1D). We observed significant differences between pre-fMRI and post-fMRI RD *(RD_individual_* : *t(19) = -8.99, p <.001 ; RD_group_ : t(19) = -9.6, p <.001)*. This result suggests that the RD between participants’ ratings and the rated social entities decreased after participants performed the fMRI task.

Following previous work[8], participants’ ratings for groups and individuals were generally consistent before and after scanning(Fig. 2C); participants were on average within ±1 of their original ratings for 75.2% (SEM=1.84 %) of group rating trials and only made large deviations from their original ratings > = 3 on 11.8% (SEM= 1.28%) of trials. For individual rating trials, they were on average within ±1 of their original ratings for 61.9% (SEM = 1.94%) and only made large deviations from their ratings >= ±3 on 21.6% (SEM= 2.56%). For self preferences, participants had high consistency in recalling their own preferences, with a 74.40% of trial consistency within 1 of their original ratings, and 5% (SEM= 1.17%) of trials displayed large deviations >= 3. When relating subjective self-confidence to self rating consistency, confidence ratings significantly correlated with rating consistency (t(19) = -5.09; p < 0.001), where higher confidence ratings yielded more consistent ratings.

During the fMRI experiment, participants made 70.9% of decisions correctly (SEM =2.98 %, *n*=20), after eliminating trials for which either answer was correct(i.e., same or equidistant ratings), while mean RT was 4.48s (SEM = 0.11). Replicating previous findings in Kaplan & Friston, 2019, there was a positive correlation between the absolute distance of the choice options from the stranger and task performance(group-level statistics: t(19) = 12.54, p < 0.001; Fig. 2B), and a negative correlation with RT (group-level statistics: t(19) = -2.65, p = 0.01). Notably, we identified a negative correlation between RT and accuracy (group-level statistics:t(19) = -5.33, p < 0.001), indicating quicker RTs for accurate choices.

Furthermore, our investigation focused on the impact of egocentric anchoring-and-adjustment during the transformation phase, particularly in the absolute reference frame. Examining the effect of RD and entities on choice accuracy, we observed a significant interaction between RD and entities (F(2,38) = 11.30, p=0.0015, partial eta squared (η^2^p) =0.37), where significant differences on performance were observed in the medium and high levels of RD between individuals and groups (mean differences in the medium bin = 8.5%, p = 0.004; mean differences in high bin = 5.73%, p= 0.007) (Fig. 2E). These significant differences highlight the presence of a u-shaped effect for RD_group_ with fMRI task performance that wasn’t present with RD_individual_. A u-shaped relationship between RD_group_ and accuracy means that egocentric biases in the absolute reference frame hurt task performance when the self rating and the presented groups’ ratings on a given trial weren’t too similar or different(S1 Fig.). Next, we examined the influence of rating discrepancy on accuracy separately for individuals and groups. For individuals, a one-way ANOVA with three levels of RD (low, medium and high) showed no significant differences in accuracy (F(2,38)=1.08,p=0.34,η^2^p=0.054, Fig.2E). Additionally, no significant quadratic relationship was found (t(19)=−1.16,p=0.87), indicating that self versus individual rating discrepancy (RD_individual_) didn’t significantly impact task performance. In contrast, a one-way ANOVA for performance by self-group rating discrepancy (RD_group_) revealed significant differences in accuracy (F(2,38)=10.8,p=<0.001, η^2^p =0.36, Fig.2E). Moreover, a significant quadratic effect was observed (t(19)=3.39,p=0.0015, and S1 Fig.), providing further support that the relationship between RD_group_ and performance follows a nonlinear u-shaped pattern.

Exploring the effect of memory on choice accuracy, we observed a significant interaction between rated entities and RC (F(3,57) = 6.59, p < 0.001, η^2^p= 0.25). In a post-hoc analysis, we observed significant differences between individuals and groups for the very inconsistent ratings >=±3 bin (mean differences = 14.97%, p= 0.004) (Fig. 2D). Then, we examined the relationship between performance and rating consistency (including memory for individuals and groups). We observed a significant correlation between these two variables(ρ=-0.59, p=0.005). Next, we separately analyzed the correlation between accuracy and RC_group_(ρ= -0.54, p=0.013) and RC_individual_(ρ=-0.52, p= 0.017). For both entities, we observe a significant negative correlation between accuracy and memory for the rating of others. In other words, the more consistent participants rated others’ preferences, the better they performed the fMRI task.

In our behavioral analyses, we observed an intricate relationship between egocentric anchoring-and-adjustment and flexible social comparisons requiring reference frame transformations. Replicating previous behavioral findings[8], we confirmed that AD negatively relates to the difficulty of a trial and egocentric anchoring-and-adjustment is present when rating entities in the task. In a subsequent attempt to disentangle the effects of memory(RC) and egocentric anchoring-and-adjustment(RD) on transformation trials, we found that memory was better for groups’ preferences(RC_group_). Notably, memory for both entity types (RC_individual_ and RC_group_)impacted how well participants flexibly compared a stranger’s preferences to different social entities that didn’t include themselves. Additionally, egocentric anchoring-related social comparison performance effects were selective to entities in the absolute reference frame(groups), where performance was lowest on mid-level rating discrepancy bins. Notably, this effect was absent when analyzing the relationship between accuracy and egocentric anchoring to the entity in the relative reference frame(RD_individual_). Taken together, these behavioral findings point towards egocentric anchoring exerting the most influence on social comparisons in the map-like absolute reference frame during the transformation task.

### fMRI analysis RSA searchlight

To test which regions represented egocentric anchor biases and other social inference demands during the transformation fMRI task, we performed whole-brain searchlight representational similarity analysis (RSA). Neural representational dissimilarity matrices (RDMs) were computed in a sphere centered on each voxel of the brain, and their relationships to behavioral RDMs were investigated(Fig. 3A). This allowed us to determine whether the neural representation of anchor biases in both relative (self rating discrepancy with individual rating: RD_individual_) and absolute(self rating discrepancy with compared groups: RD_group_) reference frames is different after explaining variance related to other important cognitive aspects in the transformation phase. The additional cognitive predictors were the absolute distance between the stranger and choice options(AD), that allow us to control for trial difficulty, rescaling of the relative position of the stranger’s rating in relation to the individual to its absolute position on the rating scale, as well as rating consistency for groups(RC_group_) and individuals(RC_individual_).

**Figure 3.**
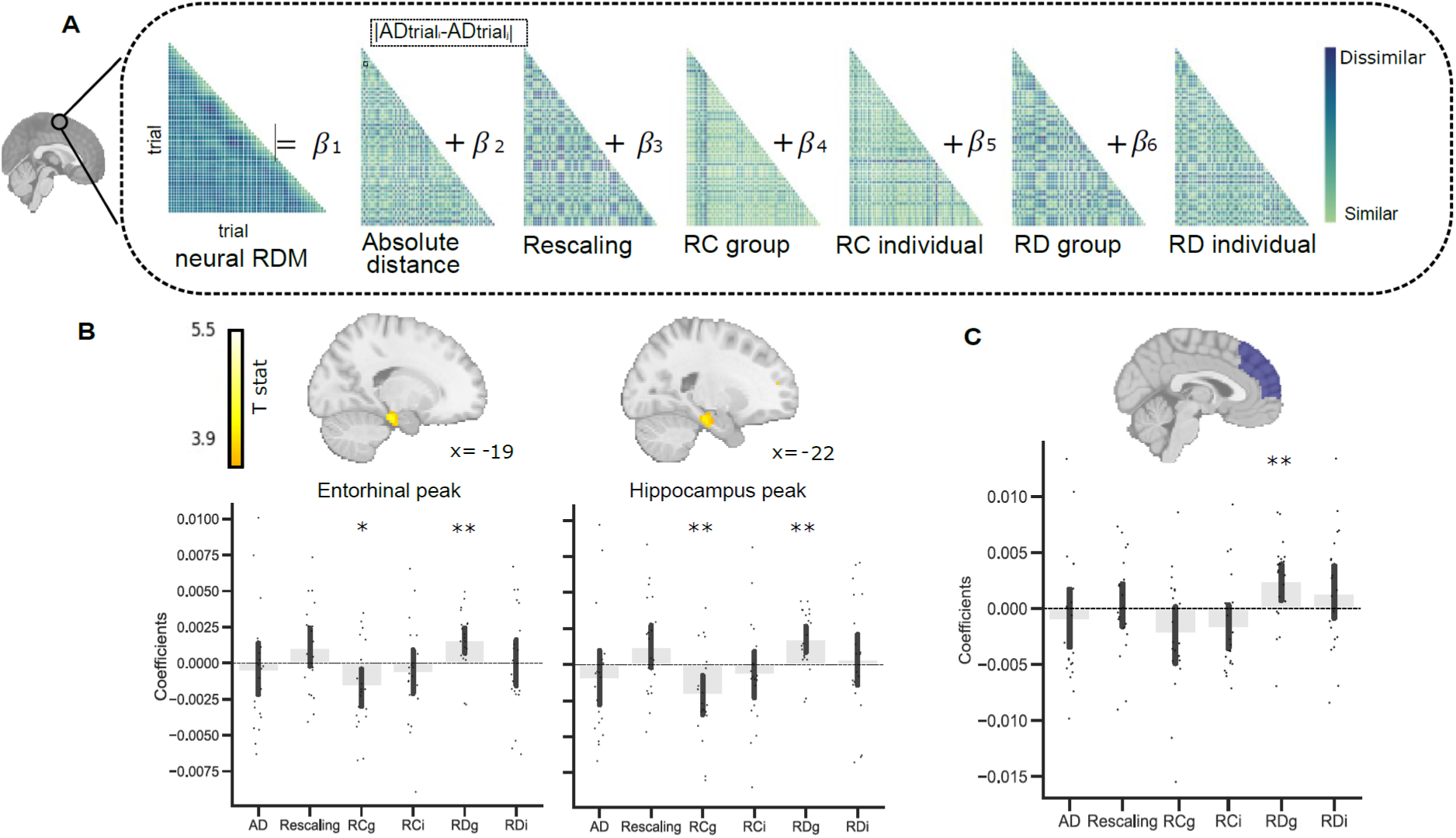
Egocentric anchoring-and-adjustment in the hippocampus. (A) Neural and behavioral representational dissimilarity matrices(RDMs) for the general linear model(GLM) searchlight analysis for one representative participant. The neural RDM was used as a dependent variable with six predictor behavioral variables. These six variables were the absolute distance between the stranger’s rating and the ratings of the groups that are being compared with the stranger(absolute distance, AD), the differences between self and others’ preferences(rating discrepancy, RD, with either groups or individuals), the cognitive demand of how far the stranger’s preference is away from the individual’s rating(rescaling), and the memory for self and others’ preferences(rating consistency, RC, for groups or individuals). To construct the behavioral matrices, absolute differences between these variables on each trial pair comparison were computed. Matrices were standardized before performing regression. (B) Top: Group activation maps for self versus group rating discrepancy(RD_group_). Differences in RD_group_ were significantly associated with left entorhinal cortex (right) (small-volume corrected family-wise error (FWE) for bilateral entorhinal cortex mask, peak-voxel corrected p=0.015) and left hippocampus pattern dissimilarity(left: small-volume corrected FWE for bilateral hippocampus mask, peak-voxel corrected p=0.046). Bottom: Barplot showing coefficients(mean ± SEM) for a 10-mm sphere around entorhinal and hippocampal peaks for each behavioral variable of interest. (C) Beta coefficients of RSA GLM in ROI of dmPFC. Bars showing beta coefficients resulting from the GLM for each behavioral variable in the dmPFC, where there is a significant dmPFC effect for RD_group_(t(19) = 2.92, p = 0.008). Each dot represents an individual participant. **= p<0.01, *=p<.05

In a whole-brain general linear model (GLM) analysis, we observed a relationship between group rating discrepancy(RD_group_) and left hippocampus pattern dissimilarity(x = –22,y = -19, z =-19; t(19)=3.78, bilateral hippocampus small-volume corrected peak-voxel p=0.046; Fig. 3B), as well as left entorhinal cortex pattern dissimilarity(x = -19,y =-25 , z=-22; t(19)=4.37, bilateral entorhinal cortex small-volume corrected peak-voxel p=0.015; Fig. 3B, S2A Fig. and S1 Table). Testing whether egocentric anchoring to the group in the hippocampus and entorhinal cortex was also present in individuals, we observed no similar effect in this hippocampal region(t(19)=-0.37, p=0.71) or entorhinal cortex(t(19)= 0.081, p=0.93) for RD_individual_. In a subsequent t-test, we tested whether this effect was specific to RD_group_, where there was no significant difference between hippocampal pattern dissimilarity for RD_group_ versus RD_individual_(t(19)=1.4, p=0.16), nor for entorhinal cortex pattern dissimilarity(t(19)=1.75, p=0.09). We then tested whether the effect found in the hippocampus peak and entorhinal peak was exclusive to the RD_group_ predictor. Using a 10 mm sphere, we examined in a post-hoc ROI analysis if this effect was present in the beta maps of other predictors. We found a significant effect for RC_group_ in both the hippocampus peak(t(19)=-3.01, p=0.007) and the entorhinal peak(t(19)= -2.49, p=0.02), but not for the other predictors(Fig.3B). A HF representation of RC is consistent with its putative role in mnemonic function[3].

Given our previous hypothesis about the role of dmPFC in egocentric anchoring-and-adjustment, we investigated whether there were any significant effects there. Despite not finding a significant dmPFC cluster for representing any variable of interest, we checked whether any effects were present in an ROI of the dmPFC. In the ROI, we observed a significant effect for RD_group_(t(19) = 2.92, p = 0.008, Fig. 3C). There were no significant dmPFC results for the other predictors. Testing whether the dmPFC rating discrepancy effects were selective to RD_group_, a subsequent t-test revealed that there was no significant difference between dmPFC RD_group_ and RD_individual_ pattern dissimilarity(t(19) = 0.74, p = 0.46). Checking whether any other brain region significantly represented any variables in our GLM, we observed a significant peak for RD_group_ in the left ventrolateral prefrontal cortex(x= -55, y=21 ,z=3; t(19)=5.55, peak-level FWE p= 0.039, S2A Fig. and S1 Table related to Fig.3B) and for RC_group_ in left fusiform gyrus(x=-34; y=-46; z=-12; cluster-level FWE p=.02; S1B Fig and S2 Table related to Fig.3B), but no other significant effects for any variables anywhere else in the brain.

To rule out that the HF, dmPFC, or anywhere else in the brain represented the self as the center of the 0-10 absolute preference rating scale on each transformation trial, we conducted a control RSA. The control analysis consisted of a whole-brain RSA searchlight informed by the proportion of ‘self recentering’(i.e., the amount of recentering needed to move a self preference rating to the center of the rating scale) on each trial. Specifically, we calculated self recentering as the absolute difference between the self rating and 5(i.e., the normal entity’s rating on every trial) divided by the absolute difference between the self rating and the furthest extreme of the scale(0 or 10) from the self rating. Crucially, we didn’t observe any significant RSA effect related to self recentering in the hippocampus peak(t(19)=1.22;p=0.23), entorhinal peak(t(19)=0.94=;p=0.35), dmPFC ROI(t(19)=0.47;p=0.64), or anywhere else in the brain.

### Pattern dissimilarity and RD_group_

We then tested whether the behavioral influence of egocentric anchoring on our task was also present in HF and dmPFC representations. To investigate whether pattern dissimilarity in the HF and dmPFC followed the u-shaped effect found in the behavioral results between RD_group_ and Accuracy (Fig.2E and S1 Fig.), we conducted quadratic regression analyses. These analyses let us determine if there was a linear or quadratic effect between pattern dissimilarity and RD_group_. However, we didn’t find a significant u-shape effect in any region examined (hippocampus peak t(19)= -0.84, p=0.79, entorhinal peak t(19)= -1.42, p=0.91 and dmPFC ( t(19) = -1.26, p = 0.88). We also investigated the possibility of a linear effect between pattern dissimilarity and RD_group_. However, no significant linear effects were found in the hippocampal peak(t(19)= -0.66, p=0.74), entorhinal peak (t(19)=-1.65, p=0.94), and the dmPFC ROI(t(19) = -0.44, p = 0.67). Additionally, we investigated whether participants who changed their pre- to post-fMRI ratings closer to their own preferences(increased egocentric anchoring) exhibited different neural representations compared to those who exhibited less of this distortion. However, we didn’t observe any significant individual differences in neural representations in the hippocampal peak((rho=-0.29;p=0.21), entorhinal peak(rho=-0.21; p=0.38), or dmPFC(rho=-0.23; p=0.33) related to increased egocentric anchoring over the course of the task.

### RSA social entity model comparison

To examine whether the hippocampus, entorhinal cortex, and dmPFC preferentially represented group identity, individual identity, or trial congruence during transformation trials, we generated trial-by-trial correlation matrices and contrasted them with the three aforementioned conceptual matrices. A trial was considered congruent when the individual and the social group involved in the transformation trial generally aligned to the same social category(i.e., an individual with an urban or rural background being paired with a respective city or village dweller group in the condition). This analysis allowed us to explore the impact of various task features on the regions’ pattern dissimilarity. We hypothesized that trials featuring the same groups would exhibit greater similarity in their hippocampal representation than trials featuring the same individuals or congruent individual-group entities. For all of the regions studied, we observe significant differences in model fits (hippocampus peak, F(2,38)= 29.08 , p<0.001; entorhinal peak, F(2,38)=15.04 ,p<0.001 and dmPFC F(2,38)=12.88,p<0.001). Following a significant ANOVA for each brain region, a paired T-test was performed to test pairwise differences between the three models. In the hippocampus peak, there was a significant difference between the group vs congruence models(t(19)=5.46, p<0.001) and group vs individual models(t(19)=6.37, p<0.001). Lastly, for the entorhinal peak, we observed a significant difference between group versus congruence models(t(19)=2.30, p=0.03) and group versus individual models(t(19)=4.68, p<0.001). These results demonstrate a consistent pattern across the regions, where the group model consistently outperformed both the individual and consistency models, suggesting that the group model better captures the underlying neural representations. In slight contrast for the dmPFC, there was no significant difference between the group vs congruence models(t(19)=1.97, p=0.06), but there was still a significant difference between group vs individual models(t(19)=5.15, p<0.001). The negative results for the individual model could be interpreted as HF and dmPFC representations being more similar when comparing different individuals, as opposed to the same individuals. Still, the possibility remains that the negative results for the individual identity model could be a byproduct of representational overlap with the group model when a condition for a trial has both the same individual and group identity.

We then ran a searchlight RSA to test if there were other brain regions that were significantly modulated by the difference between the three models. We observed two significant peaks: one in the thalamus (x=-7,y= -16,z= 9; F(2,38)= 13.42,pFWE<0.001, S3 Fig. and S3 Table) and another in the ventral striatum(x=27, y=18,z= 6; F(2,38)= 12.38, pFWE<0.001, S3 Fig. and S3 Table). For the thalamus, we observed significant differences for the individual versus congruence model(t(19)= 3.88,p<0.001) and group versus congruence model(p<0.001, t(19)=3.63), while not finding any significant difference between individual and group models(t(19)=-0.12, p=0.9). For the ventral striatum peak, we observed significant differences between the group versus individual models(t(19)=-2.58, p=0.018), individual versus congruent model(t(19)=3.35, p=0.0034), and group versus congruence models(t(19)=4.25, p=0.002).

In sum, we tested whether neural representations of implicit egocentric anchoring biases are present in the dmPFC, HF, or anywhere else in the brain when making social inferences that require flexibly comparing a stranger’s preference to multiple social entities in a map-like absolute reference frame. We found that the HF including the entorhinal cortex, the dmPFC, and ventrolateral PFC maintained representations of preferences in the absolute reference frame that were implicitly anchored to self preferences. Additionally, HF representations signaled how well preferences in the absolute reference frame were remembered, where preferences that were better remembered were represented more distinctly. Lastly, the HF and dmPFC were found to prioritize the identity of social preferences in the map-like reference frame.

## Discussion

We tested whether map-like representations of abstract social knowledge are susceptible to egocentric anchoring-and-adjustment. Replicating findings from social anchoring[25,26], we observed egocentric anchoring-and-adjustment when participants rated entities(Fig. 2A) and map-like social inference behavioral performance effects from the Kaplan & Friston paradigm(Fig. 2B). Highlighting the presence of egocentric anchoring in the HF and dmPFC, hippocampal-entorhinal and dmPFC pattern similarity was related to self versus group rating discrepancy(Fig. 3). Notably, the hippocampus and entorhinal cortex also represented how well group preferences were remembered, which was a key indicator of task performance(Fig. 3). Testing whether the HF and dmPFC preferentially represent group identity, individual identity, or trial congruence during transformation trials, we find that the HF selectively represents group identity(Fig.4). In what follows, we speculate on how egocentric anchoring potentially shapes abstract knowledge codes and relate our findings to the wider literature.

**Figure 4.**
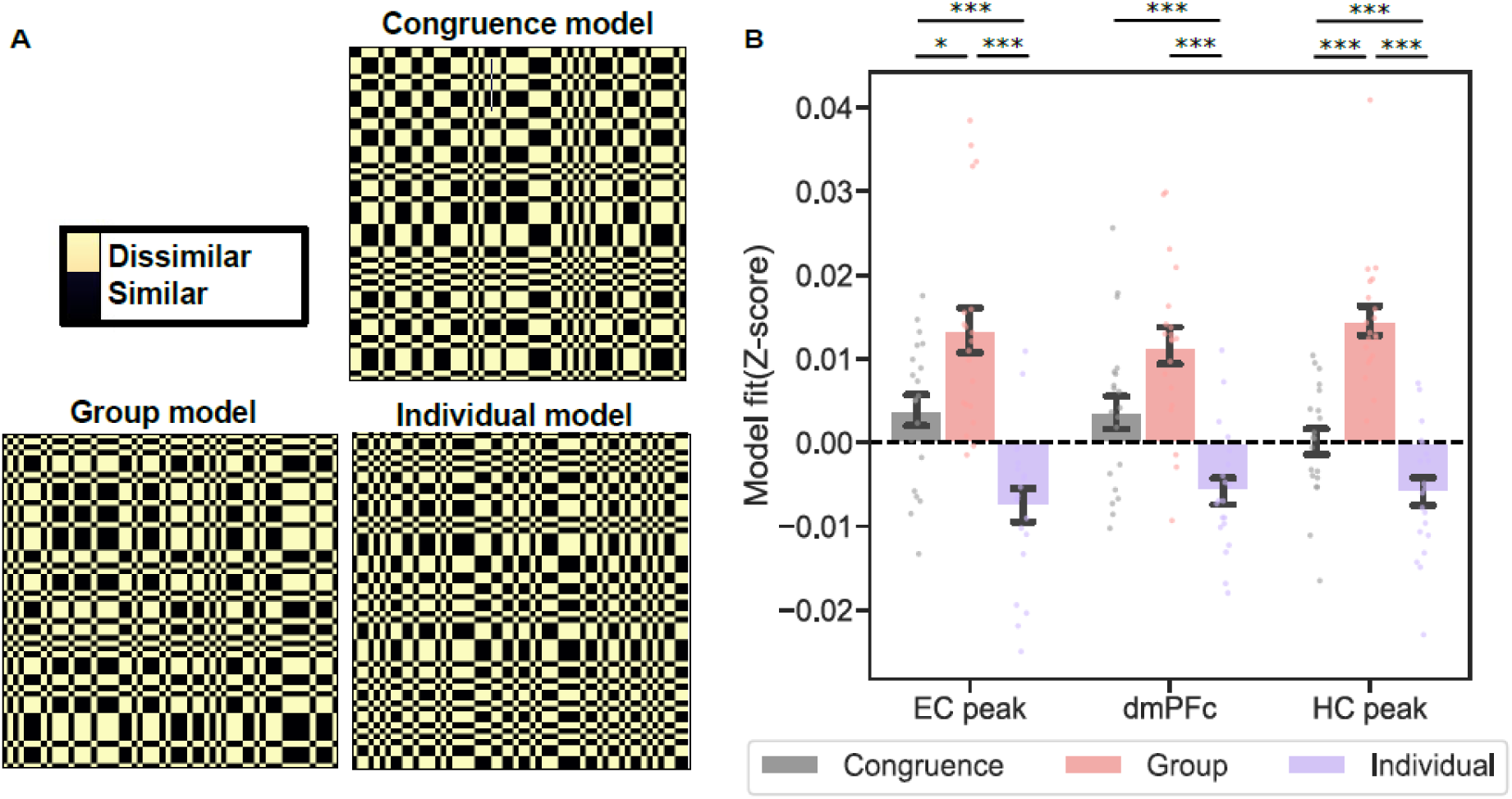
Neural representational model comparisons. (A) Model matrices based on hypothesized representational conditions: congruent versus incongruent individual and group entities(congruence), group trial identity(group), and individual trial identity(individual). (B) Correlation between model RDMs and neural RDM in the left hippocampus peak(HC), left entorhinal peak(EC), and dmPFC, where group trial identity is the winning model. All error bars showing mean ± SEM with dots representing individual participants. ***= p<0.01, *=p<.05.

### The generality of egocentric biases on allocentric coding

Our experiment doesn’t test whether HF and dmPFC representations of egocentric anchor biases during map-like inferences are selective to the social domain [12], or are more generally applicable to abstract knowledge [28]. Comparable work has uncovered the co-existence of egocentric and allocentric representations in different brain structures for abstract knowledge[29], which allows for the possibility that these parallel representations could be flexibility modulated depending on task demands. The switch between egocentric and allocentric processing could be flexibly adapted depending on task demands, which is supported by recent evidence showing personal biases can facilitate emotion prediction [30] and learning more generally[31]. Future work can study the relationship between egocentric anchoring and hippocampal generalization to uncover when self-projection is beneficial or detrimental in spatial and non-spatial inference tasks[21,32]. Despite observing significant egocentric anchoring representations in the HF and dmPFC, we can’t confirm that this subjective computation occurs on every trial across all participants. Future work with larger samples can more closely examine how individual differences in egocentric anchoring-and-adjustment changes HF and dmPFC representations during flexible social comparisons.

### Effects of social cognition and relationship with the hippocampal formation

Our findings add to a growing literature highlighting the importance of the HF in assimilating social knowledge[7–10,33,34], although an important caveat to these findings is that no study has demonstrated they are specific to cognitive map-like or non-isometric/cartesian representations of social knowledge. Further investigation of the effect of subjective biases on cognitive maps of social space[7,9,10] and assimilation of knowledge in social networks[35–38] could yield insights into whether there are differences in subjective and objective map-like coding. Moreover, investigating subjective confidence and its relationship to egocentric anchoring in the HF could be a promising avenue of investigation. Lastly, we don’t specifically investigate spatial or social perspective switching computations that likely rely on regions like the retrosplenial cortex(RSc)[13]. Future work can address whether social and spatial perspective transformations require similar underlying computations and how they might differ in areas like dmPFC or RSc [15,39].

### The specificity of anchoring to groups versus individuals in allocentric reference frames

For neural representations of rating discrepancy(egocentric anchoring) and consistency(i.e., memory strength), we find that our ROIs represent these behavioral variables for group, but not individual ratings. Moreover, these RSA effects are paralleled by behavioral performance results(Fig.2D-E), where the accuracy of social inferences is most strongly determined by self-group rating discrepancy and group rating consistency. Taken together, these results suggest that the most crucial aspects of flexible map-like social inferences are how well attributes of the social groups in our task are learned and remembered in the absolute reference frame. An important caveat to these HF and dmPFC findings is that we cannot determine whether our effect is specific to egocentric anchoring for groups, or more generally relevant to any social entity in the absolute reference frame. This is especially important because groups or any collective minded thinking can be seen as more allocentric[40], independent of any explicit reference frame transformation. Moreover, rating consistency was much lower for individuals than groups. This was likely because the groups, city and village dwellers, were more familiar than specific fictive individuals to participants. To resolve this issue, future studies can alternate whether individuals or societal groups are in the initial relative position to the stranger in our task and test how that changes task performance. Notably, different individuals were interchangeably used as the initial reference point and choice options in Kaplan and Friston(2019), which had similar behavioral results as this study. Therefore, we predict that our fMRI results would hold if the stranger was learned relative to a group and subsequently compared to individuals in the absolute reference frame.

### Relating dmPFC effects to the Social Anchoring Literature

Building on previous work by Tamir and Mitchell (2010), we observed a relationship between dmPFC representational similarity and social anchoring(self versus group rating discrepancy) in our task. Previous work isolated dmPFC fMRI signals that related to self-other discrepancy when inferring mental states of one individual[27]. In contrast, our dmPFC self-other discrepancy results extend to nonlinear social inferences that involves learning a stranger’s preference relative to an individual and then extrapolating whether the stranger’s preference is more similar to one of two groups’ preferences. Consequently, we show that dmPFC isn’t limited to its putative role of guiding one to one self-other comparisons[27,41] and can extend to more complex social inference of relating multiple social entities to each other.

### dmPFC versus Hippocampal-Entorhinal Contributions to Social Learning

Despite the dmPFC and HF being commonly linked to social [27,41] and spatial [19] learning respectively, we find that both regions represent egocentric anchor biases during social inference. One of the key differences between the dmPFC and hippocampal-entorhnal system in our task is that the HF also represented how consistently preferences were rated. Finding hippocampal-entorhinal representations of how well groups’ preferences are remembered is consistent with the region’s putative role in long-term memory[3]. Additionally, we find that the HF selectively represents group identity during transformation trials. In contrast, we observed that the dmPFC represented group identity over individual identity, but not individual-group identity congruence. These results are consistent with the idea that the HF maintains a map-like representation of one’s social network[15], while dmPFC has a more general role in integrating multiple sources of information[42]. Taken together with work showing that knowledge storage of these self preference ratings relies on ventral mPFC[43], our results imply that dmPFC is ideally positioned in the brain to interface self-knowledge with map-like knowledge of others in the hippocampal-entorhinal system.

## Conclusion

We highlight how personal preferences are present in the HF and dmPFC when making inferences about how others relate to each other. Future work can determine whether similar egocentric biases are transferable to cognitive map-like representations in spatial and other non-spatial domains[17]. By further disentangling the role of the self in shaping hippocampal models of the world, a better understanding of variability in allocentric representations will be formed.

## Methods

### Ethics Statement

Study participants were compensated and gave informed written consent to participate. The study was approved by the local research ethics committee at Universitat Jaume I (ethics reference: CD/10/2022). The study was conducted in accordance with Declaration of Helsinki protocols.

### Participants

Thirty participants (16 female, with a mean age of 21.16 years, SD=3.46) took part in an experiment conducted at Universitat Jaume I, using an online recruitment system. Four participants were excluded due to poor performance (<60%). A further four participants were excluded due to excessive movement during the experiment and two participants were omitted due to technical issues during image acquisition resulting in a final sample of 20 participants. All participants were right-handed, had normal or corrected-to-normal vision, and had no history of neurological or psychiatric disorders.

### Task

Stimuli were presented using PsychoPy (v 2020.1.1) toolbox running in Python 3.9. The task comprised distinct phases: anchor phase, transformation phase fMRI task, and rating consistency phase (Fig. 1).

### Anchor phase

Participants first provided likelihood ratings for 105 everyday activities adapted from [8](listed in Supplementary Information) on a scale from 0 to 10 (0 indicating impossibility, 10 indicating extreme likelihood). Following each question, participants provided a confidence rating on a scale from 1 to 5(from no idea to very sure), reflecting their confidence regarding their own preferences. To achieve a more varied distribution of ratings, we excluded the 30 trials that were nearest to the mean of each participant’s preference, resulting in 75 trials for the subsequent phase.

Participants were then introduced to two individuals with a brief paragraph description and the name of two stereotypical groups(city dwellers and village dwellers). To make the individuals easier to learn, each fictitious individual was created to have preferences either more compatible with city or rural village living and were gender matched to the participant. Participants were given as much time as they needed to read the full description of the two individuals. Subsequently, participants judged the entities’ preferences. Using the same 0-10 scale for self rating, participants expressed what they felt was the likelihood of each entity partaking in a variety of everyday scenarios within a 10 second time limit [8]. Participants only provided preference ratings for a given scenario once before and after the fMRI task.

### Transformation fMRI-phase

Participants then performed a brief practice version of the fMRI social decision-making task(adapted from Kaplan & Friston, 2019) outside of the scanner, until they achieved a performance level of at least 70%. During the transformation task, participants inferred a stranger’s preference relative to one of the familiar individuals using their previous ratings for the individual entities. Subsequently, in a two-alternative forced choice task (2AFC) they determined whether that stranger’s preference is more similar to a normal group (always a 5), or one of the two groups they previously rated[8] within ten seconds. Participants estimated the stranger’s preference using a scenario, and a cue of less or more related to the preferences of the previously presented individual (e.g., The stranger eats spicy food more/less than the individual). If the cue was “more”, participants had to add half of the difference between the individual preference and the high extreme of the scale (10) to the individual entity’s preference. If the cue was “less”, participants had to subtracted half of the difference between individual preference and the low extreme of the scale (0 to the individual entity’s preference). In contrast with the Kaplan and Friston study that had participants compute the stranger as slightly(¼) or very(¾) less/more than the highly familiar initial reference individual[8], we limited it here to only halfway between the reference individual and the closest extreme. This change permitted us to reduce the cognitive demand for participants since they needed to remember their ratings for four distinct entities. Participants were instructed that it was a different stranger on each trial. Crucially, participants never received any feedback during the task.

We had four conditions, each based on the individual entity used to infer the stranger’s value and the group entity presented as a choice option in the 2AFC(Fig.1A). The four different conditions were used to control for different levels of similarity between the rated entities and the participant, as well as schematic congruence between individuals and groups(level of attribute similarity between an individual and group entity in a given trial). Given the different compatibility between each rated individual and group, there were two conditions with a congruent rated individual and group(e.g., the urban oriented individual with city dwellers; the rurally oriented individual with village dwellers) and two involving an incongruent rated individual and group(i.e., the rurally oriented individual with city dwellers; the urban oriented individual with village dwellers). Due to time constraints for fMRI scanning, we excluded an additional 15 scenarios from the remaining 75 scenarios for the fMRI task, specifically those where participants rated the group preferences as 5 to avoid overlap with the normal group. Participants were therefore tested on the same 60 scenarios for all four conditions during the transformation phase, resulting in a total of 240 trials. These trials were distributed over four fMRI runs, with each run consisting of 60 trials that were assigned completely randomly. Trials were self-paced, allowing participants to determine the time for each trial(mean=4.48s) with a maximum allowed RT of 10s. As in the Kaplan & Friston study[8], any transformation trial where both choice options were correct wasn’t included in the analyses All of the information needed to perform the task was presented at once on the screen, and the aspects of the task that varied from trial to trial were the egocentric reference frame(individual), the cue (that connects individuals preferences and stranger preferences), and the choice options(see Fig. 1A). Participants had a maximum of 10s to make a decision, followed by a jittered ITI period (mean = 2.5 s; range = 1-4s), where a white fixation cross on gray background was presented. After completing the fMRI task, participants were asked to give ratings for themselves and the four entities again to test the consistency/memory for their preferences.

### Anchoring-and-adjustment analysis

To study the anchoring and adjustment effect during the anchor phase, we applied a linear mixed effect model(equation 1). Our focus was on studying the relationship between participant RTs and rating discrepancy(RD), calculated as the absolute differences between participants’ self-preferences and others’ preferences, when inferring others’ preferences. To assess differences between egocentric anchoring and adjustment for individuals and groups, we included the anchor entity and the interaction with RT as predictive variables in the model.

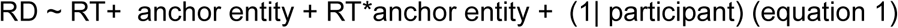

We tested for egocentric anchoring-and-adjustment by incorporating the fixed effect of RT, which is the positive correlation between RT and self–other discrepancy [25,26]. We added participants as nested variables in the model. Trials in which participants did not provide a response within the 10-second window during the self-rating or anchor phase were excluded from the analysis. This exclusion was necessary due to the impossibility of calculating the rating discrepancy variable in such instances. RTs were log-transformed to address their right-skewed distribution, and we excluded responses whose log RTs were >=2.5 SD from the grand mean[44].

### fMRI acquisition

Whole-brain structural and functional MRI data were acquired using 3T General Electric Signa Architect magnetic resonance imaging (MRI) scanner using a 32-channel head coil. Four whole-brain functional runs were acquired using a multi-band multi-echo (MBME) EPI sequence (voxel size= 3 mm isotropic, TR = 2 s; TEs = 18.5, 34.16, and 49.82 ms,; flip angle θ = 80; resolution matrix = 80 × 80, and a multi-band acceleration factor = 2). We used a functional sequence that sets the slice angle of 30° relative to the anterior-posterior commissure line to minimize signal loss in the medial temporal and orbitofrontal regions [45]. Structural T1-weighted images were acquired using an MPRAGE sequence (TR = 2.5 s; flip angle = 8, and slice thickness = 1 mm).

### Beta series modeling

Pre-processed BOLD time series data was analyzed using a Least Squares-All (LSA) general linear model (GLM) that afforded estimation of the specific activation patterns elicited by each trial and run. The model includes every single trial as a regressor, and various confound regressors[46] composed of 6 motion regressors, 6 physiological noise regressors, and a global signal regressor. The last regressor was included to help control for global effects or potential confounding factors that might affect the entire fMRI brain signal[47]. Analysis was performed using Nilearn v 0.10.1, a Python library for statistical learning on neuroimaging data (https://nilearn.github.io).

### fMRI data analysis

#### Calculation of behavioral dissimilarity matrices

For each participant and run, we calculated trial by trial dissimilarity matrices for each of the behavioral variables of interest: absolute distance between choices(AD), rescaling, self versus individual rating consistency(RC_individual_), self versus group rating consistency(RC_group_), self versus group rating discrepancy(RD_group_), and self versus individual rating discrepancy(RD_individual_). These matrices were constructed by computing the absolute differences between the values of these variables across trials.

### Representational similarity analysis (RSA)

We performed a whole-brain searchlight-based multiple regression RSA to examine brain regions in which changes in dissimilarity between neural activity patterns in the trials is explained by the behavioral variables of interest. To create the neural representation dissimilarity matrix (nRDM) for each voxel, a spherical searchlight was run by defining a sphere with a radius of four voxels that was moved across the brain. For each sphere, we extract the beta coefficients for the voxels within that sphere, which are derived from the LSA-GLM model (as detailed in the Beta-Series Modelling section). This extraction was repeated across all trials, producing a vector of beta coefficients for each sphere location [48]. Next, we perform a single value decomposition analysis to reduce the dimensionality to ten dimensions, where these first ten dimensions capture 76.04% variance for all participants and searchlights. These reduced vectors are then correlated to generate an nRDM, with dimensions determined by the number of trials in the transformation phase. This neural RDM is then used as the dependent variable in the model for each voxel. To investigate how changes in behavioral variables relate to neural fMRI signal patterns in different brain regions, we applied a voxel-wise GLM to predict the nRDM with six behavioral predictors (Fig. 3A). Behavioral matrices were standardized before performing regressions. For each participant, the multiple regression-based RSA provided a beta map for each of the six predictors. These beta maps indicate the relative influence of each predictor on the dissimilarity of neural activity patterns within the searchlight spheres. In a second-level analysis, the resulting beta estimates for each participant and for each voxel were testing the regression coefficients against zero using one sample t-test.

In addition to the searchlight-based RSA, we ran a ROI-based RSA in dmPFC. We computed the trial by trial nRDM, extracting the beta coefficients for all voxels within the ROI and computed correlations between all transformation trials. We also applied the aforementioned dimensionality reduction. The resulting nRDM was then introduced as the independent variable in the same GLM(Fig. 3A) employed for the searchlight analysis.

To control for the presence of any type of self recentering representation, we ran a control whole-brain searchlight RSA. This analysis was informed by the proportion of ‘self recentering’(i.e., the amount of recentering needed to move a self preference rating to the center of the rating scale) on each trial. We also checked for the presence of these effects in 10mm ROIs around our hippocampal and entorhinal RD_group_ peaks, as well as in the dmPFC ROI.

### Pattern dissimilarity by RD_group_

We extracted the beta values within a 10mm radius sphere around the hippocampal peak, entorhinal peak, and dmPFC ROI for each participant from the corresponding beta image for each trial. Subsequently, we concatenated all trials from different runs and constructed correlation matrices for trials that share the same RD_group_ value, resulting in nine representational dissimilarity matrices. Trials with a singular representation for an RD_group_ category were excluded from the analysis. To quantify dissimilarity, we convert correlation values to dissimilarity using the 1 - correlation. Following this transformation, we calculated the mean pattern dissimilarity value for each nRDM.

To study the relationship between accuracy and pattern dissimilarity with RD_group_, we calculate two regression models for each participant. We add a quadratic term 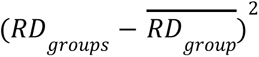 to account for potential symmetrical quadratic relationships, where the β2 coefficient expresses the strength of the quadratic relationship between dependent variables and RD_group_ .

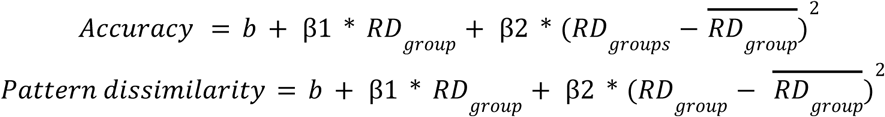

For the group-level analysis, we test if the β2 coefficients were significantly different from zero, which allows us to study the quadratic relationship. To study the relationship between these two models, we correlated the beta values for each regression between them. This correlation between coefficients across participants helped us identify consistent patterns in the direction of the quadratic effects.

### Model matrix comparisons

To test specific hypotheses about how groups, individuals, and trial congruency are represented, we compared trial by trial correlation matrices for pattern similarity data to model matrices(Fig. 4A). Beta images for each trial, resulting from the preprocessing step in the *Beta series modeling* section, were masked with a 10 mm radius spherical mask on significant peaks identified in the RSA searchlight analysis and in the dmPFC ROI. The beta values underwent pairwise comparisons resulting in a trial by trial nRDM, for each participant and run. We computed three model matrices: (1) congruence model, (2) group model, and (3) individual model.

For the congruence model, transformation trials were classified as congruent when the egocentric reference frames (individuals) and allocentric reference frames (groups) belong to the same social category (i.e. trials involving city individual and city group). Conversely, trials were considered incongruent when the egocentric and allocentric reference frames belong to different categories(i.e., city individual and rural people group). To calculate congruence matrices, we hypothesize that trials within the same congruence condition will exhibit higher correlations with other trials sharing the same condition. To calculate the group model matrix and the individual model matrix, we hypothesized that trials that involve the same social groups or the same individuals will contain more similar representations. Model matrices were constructed assigning a value of 0 to trials with a hypothesized high correlation (similarity), and a value of 2 to those with a hypothesized low correlation(dissimilarity)(S4 Fig. related to Fig. 4).

To evaluate model fit, we computed a partial correlation between participant average trial-wise pattern similarity data and each of the 3 model matrices using the lower half of the matrix (excluding comparisons involving the same trials) after eliminating the effect of condition. At the group-level, we computed one-way ANOVAs and a paired T-test. These statistical analyses allowed us to compare fits across models.

In addition to the model matrix comparison RSA, we performed a whole-brain searchlight-based multiple regression RSA to examine the brain areas in which the change in dissimilarity between neural activity patterns in the trials is explained by the different three models. Subsequently, we performed an ANOVA to compare the difference between the coefficients for each model and a paired t-test to determine which model was driving the observed effects.

### fMRI statistical analysis

For the whole-brain results, the statistical analyses are performed using 10 mm radius spheres in Nilearn toolbox (v.0.10.2)[49] around the respective peak-voxel specified in the voxel-wise GLM analysis. We report peak-voxels in the hippocampus, dmPFC and entorhinal cortex that survive small-volume correction for multiple comparisons (p<0.05) based on bilateral hippocampal, dmPFC, and entorhinal cortex masks. We defined the bilateral hippocampal region using the Wake Forest University (WFU) PickAtlas, integrated with SPM [49,50]. The entorhinal cortex mask was created based on prior literature [8,52,53]. For the regions of interest (ROI) analysis, we define the dmPFC mask [54] as the medial wall of the prefrontal cortex that forms part of Brodmann areas 9 and 10. Additionally, medial parts of Brodmann area 8 adjacent to area 9, as well as the dorsomedial portion of area 32, have been included. For all analyses outside the regions of interest (ROIs), we report activations surviving an uncorrected statistical threshold of p=0.001 and correction for multiple comparisons at the whole-brain level (FWE p<0.05) either at the peak-voxel or cluster-level. Coordinates of brain regions are reported in MNI space. Analysis was performed using a Python open-source RSAtoolbox v.3.0 (https://github.com/rsagroup/rsatoolbox, [55]) and statsmodels toolbox v. 0.14.0 (https://github.com/statsmodels/statsmodels, [56]).

## Acknowledgments

We thank María Baena Pérez and Esteban Villar Rodríguez for assisting with fMRI piloting, Maria Picó Pérez for fMRI preprocessing advice, and Ameer Ghouse for statistical and programming assistance. We thank Shahar Arzy and Maya Visser for feedback on a previous version of this manuscript. We thank General Electric for providing a pilot version of their multi-echo fMRI sequence package. This research is supported by grants awarded to RK from the Valencian Community’s Program for the Support of Talented Researchers (CIDEGENT/2021/027), Universitat Jaume I Research Advancement Plan(UJI-B2022-45), and Spanish Science, Innovation, and University Ministry(PID2021-122338NA-I00).

