## Supplementary Information for "Egocentric anchoring-and-adjustment of social knowledge in the hippocampal formation"

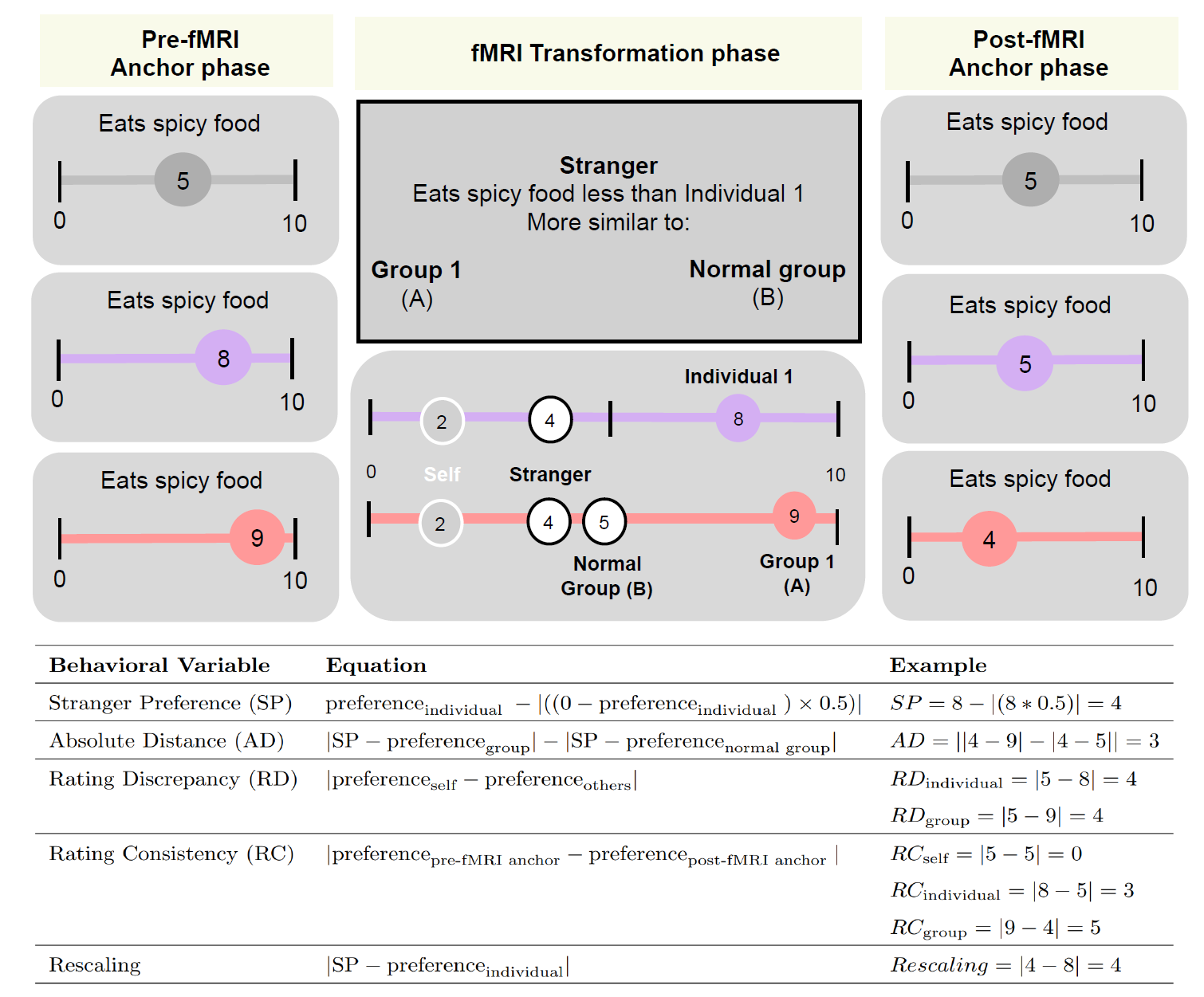

**Supplementary Figure 1:** Example fMRI trial where participants´ pre- and post-fMRI task ratings for themselves(the rating in gray) and the other relevant entities(individuals in purple, groups in pink) are highlighted. Ratings for the self aren't needed to accurately perform the fMRI task. Presented below in the table are how the experimental variables of interest would be calculated for the given preference ratings in this example trial. Stranger preferences(SP) are only used to inform the other behavioral variables and aren't analyzed in isolation in this study. Crucially, post-fMRI task ratings only are used for the Rating Consistency variables.

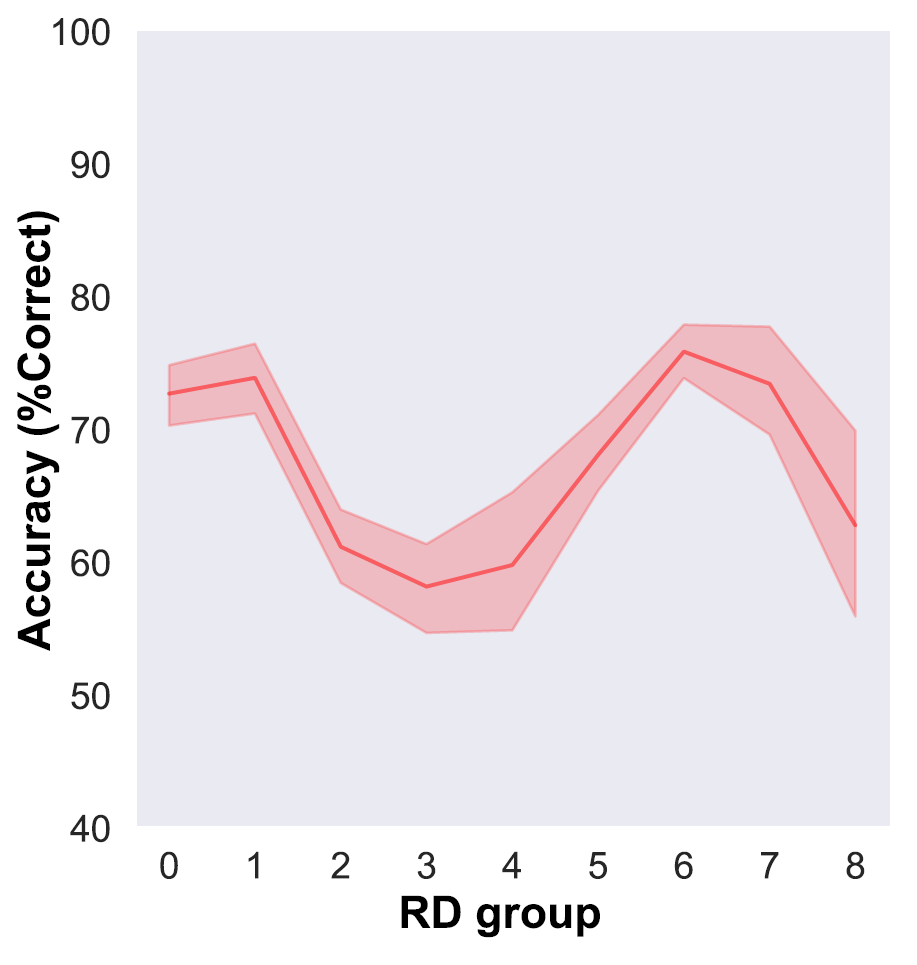

**Supplementary Figure 2:** Significant quadratic effect between accuracy and RD_group_ (t(19)=3.39, p=0.0015).

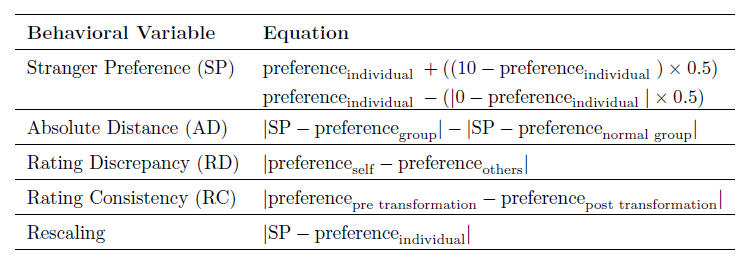

**Supplementary Table 1. Experimental variables of interest**: Stranger preferences(SP): If the cue was “more”, participants had to add half of the difference between the individual preference and the high extreme of the scale(10) to the individual entity's preference. If the cue was “less”, participants needed to subtract half of the difference between individual preference and the low extreme of the scale(0 to the individual entity's preference). Absolute distance (AD) between a stranger’s rating for a preference and the two choice options. Rating discrepancy (RD): distance between self and either other individual(RD_individual_) or group(RD_group_) preferences for a particular scenario; Rating Consistency(RC): differences between anchor phase ratings pre- and post-fMRI scanning for the self, groups(RC_group_), or individuals(RC_individual_); Rescaling: the cognitive demand of how far the stranger’s preference is away from the individual’s rating.

*Multi-echo data fMRI pre-processing*

This study used a multiband multi-echo (MBME) scanning sequence. We used TEDANA to combine the images [54-57]. Before images were combined, preprocessing was performed using fMRIPrep 22.1.1 [58]; RRID:SCR_016216), which is based on Nipype 1.8.5([46, RRID:SCR_002502). Outputs of individual echoes after slice-timing correction, head-motion correction, and susceptibility distortion correction were introduced into the TEDANA pipeline to denoise data and combine the images. To normalize scanner-space TEDANA denoised data into MNI space, we used ANTS’s antsApplyTransforms tool. For more details on the fMRIprep pipeline see https://fmriprep.readthedocs.io/en/stable/workflows.html.

*Beta series modeling*

Pre-processed BOLD time series data underwent analysis using a Least Squares-All (LSA) general linear model (GLM) that allowed to estimate the specific activation patterns elicited by each trial and run. The model includes every single trial as a regressor, and various confound regressors [59] composed of 6 motion regressors, 6 physiological noise regressors, and a global signal regressor. The last regressor was included to help control for global effects or potential confounding factors that might affect the entire brain 60]. Analysis was performed using Nilearn v 0.10.1, a Python library for statistical learning on neuroimaging data (<https://nilearn.github.io>).

105 Everyday Scenarios

1.Waking up early during the week

2.Taking an online class

3. Ironing clothes

4. Uploading picture to your social network

5. Playing board games

6. Devoting time to choose everyday clothing

7. Decorating their place

8. Having good sleep habits

9. Taking a dance class

10. Owning a pair of running shoes

11. Visiting a library

12. Talking on the phone

13. Talking on the phone in a foreign language

14. Using a motorbike

15. Buying clothes from a second-hand shop

16. Going to bed late

17. Going to a nightclub

18. Reading a novel

19. Taking a nap

20. Having a meeting with a co-worker of a different

nationality

21. Playing football with your friends

22. Having a second house in the same province/state

23. Ordering take away pizza

24. Communicate via email

25. Listening to commercial music

26. Cycling bike

27. Sleeping more than 7 hours a night

28. Going out to dinner with friends.

29. Drinking coffee in the morning

30. Watching a romantic movie

31. Eating sushi

32. Speaking on the phone in a foreign language

33. Riding the bus

34. Watching cartoons

35. Following politics

36. Watching TV on evening

37. Drinking tea

38. Falling asleep in the car

39. Playing padel

40. Eating brunch

41. Reading newspaper

42. Playing video games

43. Eating a kebab

44. Following fashion trends

45. Visiting family

46. Throwing a house party

47. Listening to music in a foreign language

48. Fixing things around the house

49. Chat via instant messaging

50. Eating vegetables

51. Eating paella on Sunday

52. Eating fast food

53. Downloading a movie

54. Going to a concert

55. Painting their home

56. Meeting friends at the pub

57. Wearing a wool sweater

58. Eating spicy food

59. Smoking a cigarette

60. Speaking Valencian with friends and family

61. Growing vegetables

62. Wearing a leather jacket

63. Going to a work conference

64. Eating exotic meal

65. Knowing your friend’s grandparents

66. Sending a message using mostly emojis

67. Traveling abroad

68. Spending a weekend on the sofa

69. Going to a yoga class

70. Taking a coffee break at work

71. Listening to podcast

72. Spending a lot of money in clothes

73. To use laptop to take notes

74. Drinking milk

75. Wearing boots

76. Changing jobs

77. Falling asleep in front of the television

78. Holidaying in a tropical location

79. Personally knowing your neighbors

80. Greeting strangers when you enter in a public space

81. Using a dishwasher

82. Buying a new car

83. Watching sport on TV

84. Going to the beach during the week

85. Taking the train

86. Hiring a person to clean their house

87. Cooking dinner

88. Joining a political march

89. Watching a soap opera

90. Socializing with people older than you

91. Speak via video chat

92. Going for a hike

93. Attending a musical

94. Jogging 5 miles

95. Visiting an art museum

96. Wearing a hat

97. Driving to work

98. Meditating

99. Receiving a work-related phone call.

100. Taking more than an hour to get ready in the morning

101. Catching a taxi

102. Being accustomed to traffic

103. Living close to nature

104. Buying designer clothes

105. Doing sudoku
